# LTD-inducing low frequency stimulation enhances p-Tau181 and p-Tau217 in an age-dependent manner in live rats

**DOI:** 10.1101/2022.03.28.486022

**Authors:** Yangyang Zhang, Yin Yang, Zhengtao Hu, Manyi Zhu, Shuangying Qin, Pengpeng Yu, Bo Li, Jitian Xu, Michael J. Rowan, Neng-Wei Hu

## Abstract

The progressive cognitive decline in Alzheimer’s disease (AD) patients correlates with the extent of tau pathology, in particular tau hyperphosphorylation, which is strongly age-associated. Although elevation of phosphorylated tau (p-Tau) on residues Thr181 (p-Tau181), Thr217 (p-Tau217), and Thr231 (p-Tau231) in cerebrospinal fluid or blood are recently proposed to be particularly sensitive markers of early AD, the generation of p-Tau during brain activity is poorly understood. A major form of synaptic plasticity, long-term depression (LTD), has recently been linked to the enhancement of tau phosphorylation. Here we show that low frequency stimulation (LFS), used to induce LTD, enhances p-Tau181 and p-Tau217 in an age-dependent manner in the hippocampus of live rats. In contrast, phosphorylation at residues Thr231, Ser202/Thr205, and Ser396 is less sensitive to LFS. Pharmacological antagonism of either NMDA or metabotropic glutamate 5 (mGluR5) receptors inhibits the elevation of both p-Tau181 and p-Tau217. Targeting ageing with a small molecule cognitive enhancer ISRIB (trans-isomer) prevents the enhancement of p-Tau by LFS in aged rats. Together, our data provide an *in vivo* means to uncover brain plasticity-related cellular and molecular processes of tau phosphorylation in health and ageing conditions.

## Introduction

Clinical evidence indicates that age-associated progressive cognitive decline in Alzheimer’s disease (AD) patients correlates with the extent of tau pathology, in particular the degree and nature of tau phosphorylation (Chang *et al*, 2021; Nies *et al*, 2021; Wegmann *et al*, 2019; Wesseling *et al*, 2020). Among the latter, elevation of phosphorylated tau (p-Tau) on residues of Thr181 (p-Tau181), Thr217 (p-Tau217), and Thr231 (p-Tau231) in cerebrospinal fluid (CSF) or blood were recently proposed to be particularly sensitive markers of early AD, long before the diagnosis of clinical dementia (Barthelemy *et al*, 2020a; Barthelemy *et al*, 2020b; Hansson, 2021; Karikari *et al*, 2020; O’Connor *et al*, 2020; Palmqvist *et al*, 2021; Wegmann *et al*, 2021). P-Tau in the brain and its subsequent release into CSF and blood is a dynamic process that changes during disease evolution. Although reports from various memory clinics indicate that p-Tau181, p-Tau217, and p-Tau231 distinguish AD from controls with high accuracy for very early AD diagnosis, the generation of these p-Tau species in patients’ brains, in particular learning and memory processes, is still unclear, mainly due to ethical and technical limitations.

The hippocampus is one of the areas that p-Tau first appears during Braak stage II of AD (Braak *et al*, 2006) and this region of brain is particularly vulnerable to age-related changes (Buss *et al*, 2021; Driscoll *et al*, 2003; Ianov *et al*, 2017; McKiernan & Marrone, 2017; Veldsman *et al*, 2021). Synaptic plasticity mechanisms at excitatory glutamatergic synapses in hippocampus, including those underlying memory function (Connor & Wang, 2016; Magee & Grienberger, 2020). Interestingly, recent studies indicate that LTD induction enhances tau phosphorylation at Ser396 (p-Tau396) (Kimura *et al*, 2014; Regan *et al*, 2015) and Ser202/Thr205 (p-Tau202/205) (Taylor *et al*, 2021) in hippocampus *in vitro*. It is still unknown whether the expression levels of p-Tau181, p-Tau217, and p-Tau231 can also be enhanced by physiological LTD induction. Whether or not their enhancement is more sensitive compared with other reported residues and the age-dependence of tau phosphorylation remains elusive.

Both NMDAR and metabotropic glutamate receptors are required for the induction of most forms of LTD by low frequency conditioning stimulation (LFS) (Collingridge *et al*, 2010). Recent evidence implicates a particular role for extrasynaptic NMDAR (Liu *et al*, 2013; Papouin *et al*, 2012) and metabotropic glutamate receptor subtype 5 (mGluR5) (Hu *et al*, 2014; Li *et al*, 2009; Luscher & Huber, 2010; O’Riordan *et al*, 2018a), both also involved in tau pathology (Benarroch, 2018; Sun *et al*, 2016; Tackenberg *et al*, 2013), in LTD induction.

Here we investigated whether p-Tau181, p-Tau217, p-Tau231, p-Tau202/205 and p-Tau396 are affected by the induction of hippocampal LTD by LFS at CA3 to CA1 synapses in the hippocampus of live rats (Hu *et al*., 2014; O’Riordan *et al*, 2018b; Ondrejcak *et al*, 2019). We assayed the local expression of p-Tau in different subregions of hippocampus at two different ages (2-3-months and 17-18-months). We found that electrical LFS preferentially enhanced p-Tau181 and p-Tau217 in an age-dependent manner without apparently affecting the levels of p-Tau at other residues that were investigated. Further, blocking either NMDARs or mGluR5 with their selective antagonists strongly inhibited the elevation of both p-Tau181 and p-Tau217. Finally, targeting ageing with ISRIB (trans-isomer) (Krukowski *et al*, 2020) prevented the increase of both p-Tau181 and p-Tau217 by LFS in aged rats. Our data provide an *in vivo* means to uncover brain plasticity-related cellular and molecular processes of tau phosphorylation in health and disease.

## Results

### Induction of LTD by LFS enhances p-Tau181, p-Tau217 in an age-dependent manner in live rats

Pyramidal neurons in hippocampal CA1 area is one of the fields that p-Tau first appears during Braak stage II of AD (Braak *et al*., 2006). Although LFS-triggered tau phosphorylation at Ser396 (Kimura *et al*., 2014; Regan *et al*., 2015) and Ser202/Thr205 (Taylor *et al*., 2021) in hippocampal slices has been previously reported, the effects of LTD-inducing LFS on the expression of p-Tau181, p-Tau217, and p-Tau231, recently proposed to be particularly sensitive markers of early AD, have yet to be described. Having developed stimulation protocols to reliably induce LTD at CA3 to CA1 synapses in live rats (Hu *et al*., 2014; O’Riordan *et al*., 2018a, b; Ondrejcak *et al*., 2019), we confirmed (Hu *et al*., 2014) that this protocol (LFS-900, 900 pulses at 1 Hz) triggered a robust persistent form of LTD that, like certain forms of long-term memory formation, is protein synthesis-dependent (**Figure S1**). We used the same protocol (LFS-900, 900 pulses at 1 Hz) in this study (**Figure 1a**). To determine the age-dependence of p-Tau enhancement by LFS, we performed our experiments in rats at two different ages: 2-3 months and 17-18 months. LFS depressed field EPSPs to 63.8 ± 7.8% of baseline in 2-3-month-old rats, and 62.4 ± 3.4% in 17-18-month-old rats. The magnitude of LFS-induced synaptic depression is thus comparable at both ages (**Figure 1b,c**).

**Figure 1.**
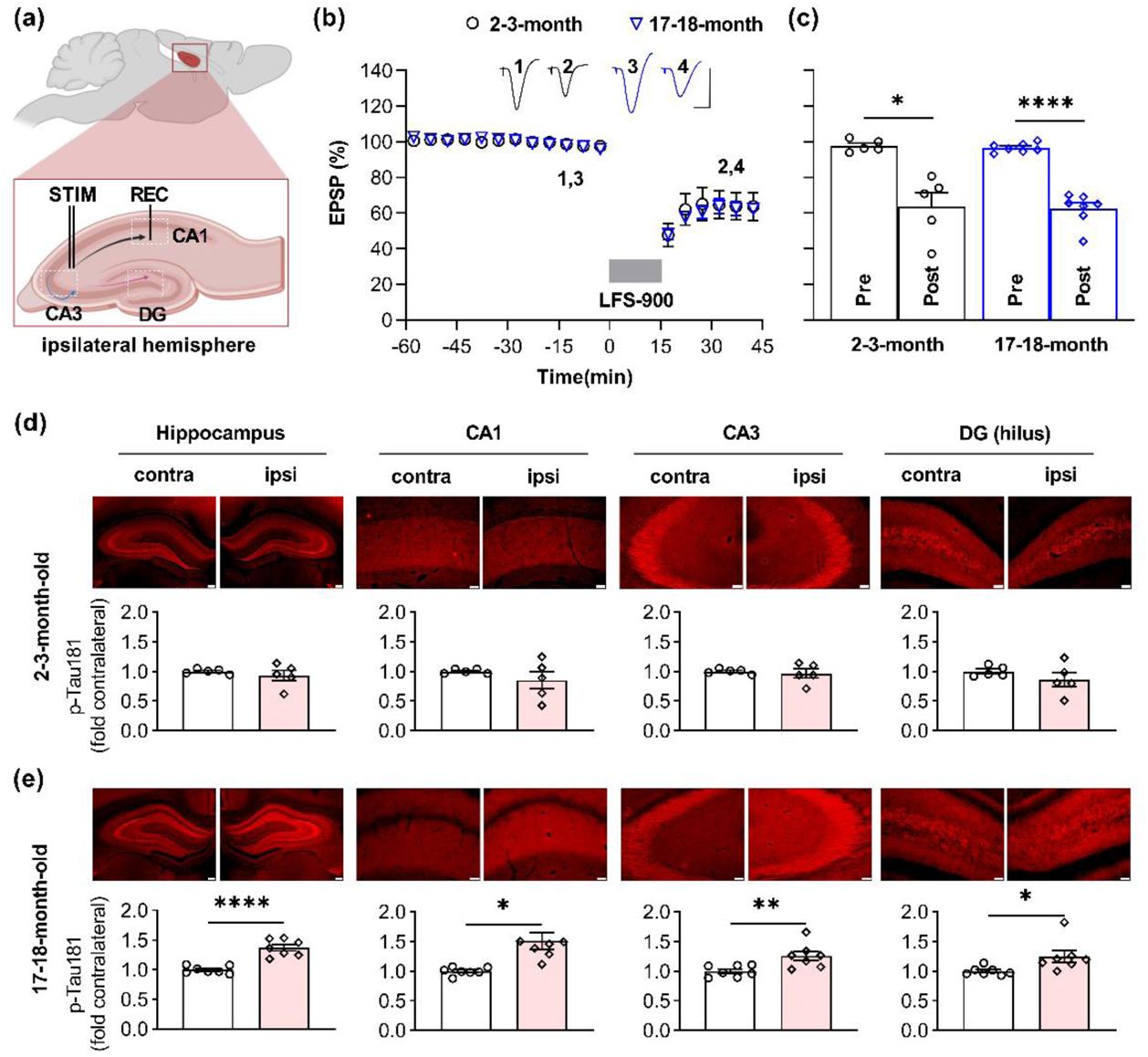
LFS promotes p-Tau181 in an age-dependent manner in live rats. (a) Schematic of the field EPSPs recording configuration in CA1 stratum radiatum (REC) overlaid with a schematic of a bipolar stimulation electrode (STIM) for Schaffer collateral axon fibers (black) in ipsilateral hemisphere. Additional excitatory projections from CA3 include local recurrent connections (blue) of CA3 pyramidal cells onto other CA3 pyramidal cells, and the back projection (pink) of CA3 pyramidal neurons to the dentate gyrus (DG). (b) Application of LFS (horizontal bar, LFS-900; 900 pulses at 1Hz) induced robust LTD at CA3-CA1 synapses in anaesthetized rats at two different ages (2-3-month, and 17-18-month). Calibration bars: vertical, 2 mV; horizontal, 10 ms. (c) Summarized EPSP amplitude 30 min post LFS. The EPSP decreased to 63.8 ± 7.8% in 2-3-month-old rats (n = 5, *P* = 0.0147 compared with Pre), and 62.4 ± 3.4% in 17-18-month-old rats (n = 7, *P* < 0.0001 compared with Pre) respectively; paired *t* test. The amplitude of LTD is comparable in both groups (two-way ANOVA, age, *F* (1, 10) = 0.09427, *P* = 0.7651). (d) The upper panel shows p-Tau181 (red) immunofluorescent staining in dorsal hippocampus (Scale bar: 200 μm), CA1, CA3, and hilus of DG (scale bars: 50 μm) from 2-3-month-old rats. The corresponding statistical results compared with contralateral side are displayed in the lower panel. The expression level of p-Tau181 was not affected by LFS in dorsal hippocampus (*P* = 0.4611), CA1 (*P* = 0.3792), CA3 (*P* = 0.6842), and DG (*P* = 0.1953); paired *t* test. (e) Immunofluorescent staining of p-Tau181 (red) in ipsilateral dorsal hippocampus, CA1, CA3, and DG from 17-18-month-old rats. LTD induction by LFS significantly enhanced the level of p-Tau181 in dorsal hippocampus (*P* < 0.0001), CA1 (*P* = 0.0220), CA3 (*P* = 0.0059), and DG (*P* = 0.0202); paired *t* test. Values are mean ± s.e.m.

The rats were sacrificed 30 min post-LFS and immunohistochemically processed for p-Tau181, p-Tau217, p-Tau231, p-Tau202/205, p-Tau396, and total tau analysis (all antibodies used are in **Table S1** and in the Methods). The expression level of total tau and p-Tau was measured in whole dorsal hippocampus, CA1, CA3, and the hilus of dentate gyrus (DG in this study) areas. The expression level in the contralateral hemisphere was used as the control. Immunofluorescent staining for antibody Tau46 or Tau5 confirmed that application of LFS did not lead to a change in total tau expression level in both age groups (**Figure S2**). Whereas no difference of p-Tau181 level was apparent in 2-3-month-old (**Figure 1d**), in 17-18-month-old animals tau phosphorylation at Thr181 was obviously enhanced in all three hippocampal fields (**Figure 1e**). We then assayed the expression level of p-Tau217 in adjacent brain slices from the same animals. Similar to p-Tau181, no difference was observed in 2-3-month-old rats (**Figure 2a**), while an enhancement of p-Tau217 was seen in CA1, CA3, and DG in 17-18-month-old rats (**Figure 2b**). In contrast, the expression levels of p-Tau231, p-Tau202/205 and p-Tau396 did not appear to change in adjacent brain slices from young (**Figure S3**) or aged rats overall, with the exception of a significant enhancement of p-Tau202/205 in the CA1 area of older rats (**Figure S4**).

**Figure 2.**
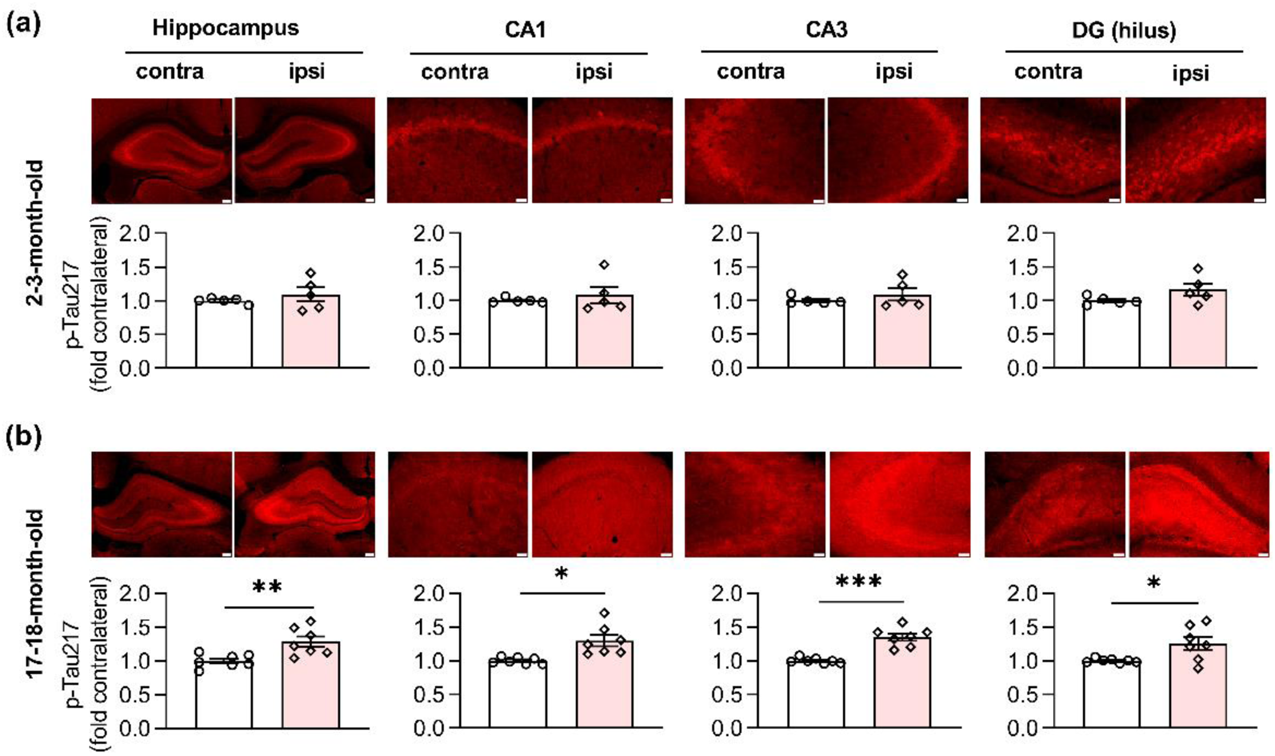
LFS promotes p-Tau217 in an age-dependent manner in live rats. (a) The upper panel shows immunofluorescent staining of p-Tau217 (red) in dorsal hippocampus (Scale bar: 200 μm), CA1, CA3, and DG (scale bars: 50 μm) from 2-3-month-old rats. The corresponding mean fluorescence intensities were summarized in the lower panel. LFS did not affect the expression level of p-Tau217 in dorsal hippocampus (*P* = 0.3820), CA1 (*P* = 0.4488), CA3 (*P* = 0.3409), and DG (*P* = 0.1567); paired *t* test. (b) Immunofluorescent staining of p-Tau217 (red) in dorsal hippocampus, CA1, CA3, and DG from 17-18-month-old rats. LFS ipsilaterally enhanced the level of p-Tau217 in dorsal hippocampus (*P* < 0.0019), CA1 (*P* = 0.0153), CA3 (*P* = 0.0003), and DG (*P* = 0.0251); paired *t* test. Values are mean ± s.e.m.

To determine if the changes in p-Tau by LTD-inducing LFS detected using immunohistochemistry referenced to the contralateral hemisphere, we measured the expression level of the same phospho-tau species using western blotting after the same conditioning LFS was applied in another cohort of 17-18-month-old rats (**Figure 3a,b**). β-actin was used to ensure equal protein loading on gels. The total tau expression level, normalized to β-actin, in either the ipsilateral or contralateral hippocampus did not differ from age-matched naïve control rats (**Figure S5**).

**Figure 3.**
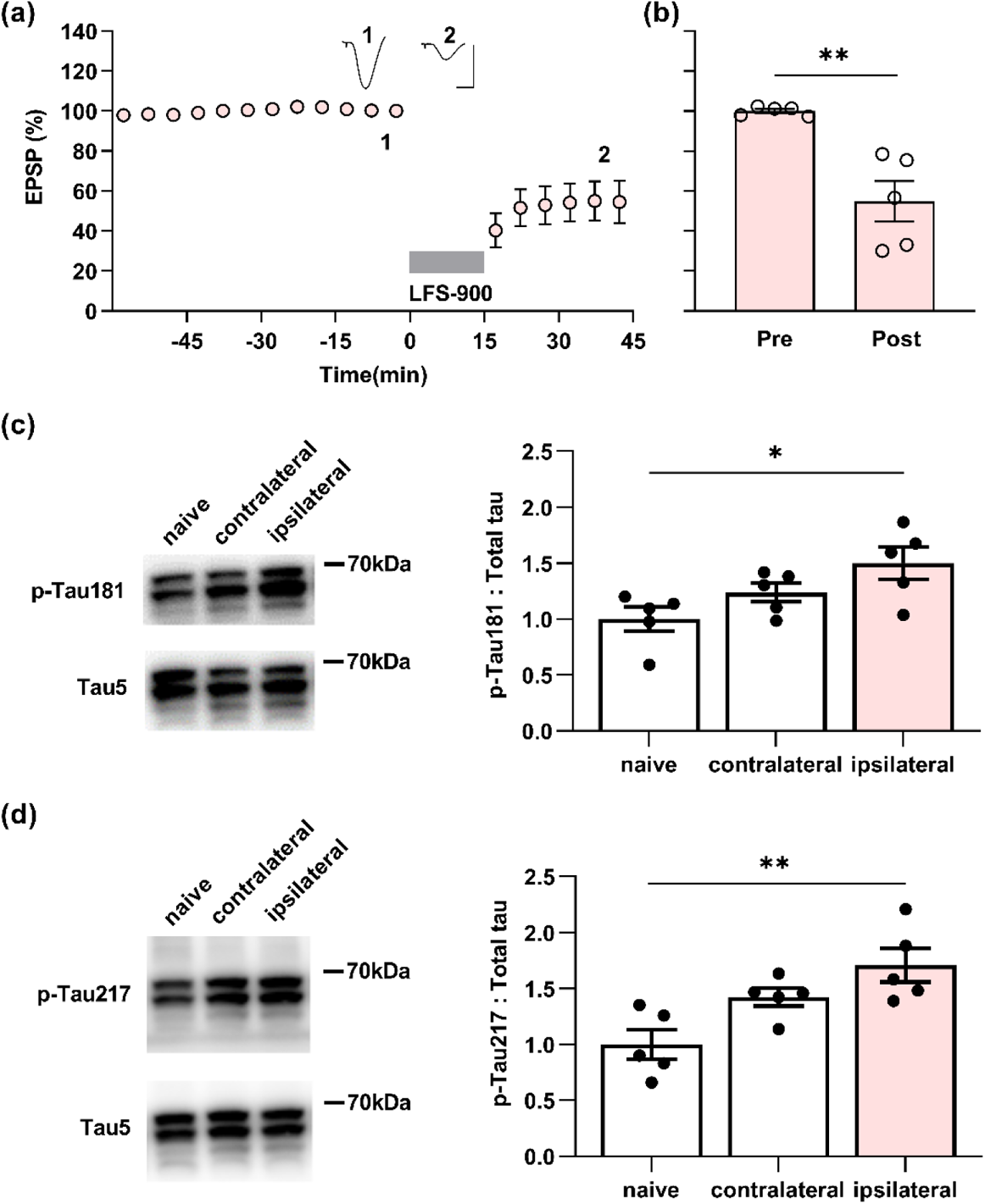
LFS enhances p-Tau181 and p-Tau217 in aged rats. (a) Application of LFS-900 induced robust LTD at CA3-CA1 synapses in anaesthetized 17-18-month-old rats. Calibration bars: vertical, 2 mV; horizontal, 10 ms. (b) Summarized EPSP amplitude 30 min post LFS. The EPSP decreased to 54.8 ± 10.2% (n = 5, *P* = 0.0110 compared with Pre, paired *t*). (c) Left panels show representative blotting band of p-Tau181 and total tau (Tau5) in the hippocampus of age-matched naïve control group and the experimental group either contralateral or ipsilateral to LFS. Statistical results of p-Tau181 over total tau was quantified and normalized to naïve control (n = 5 per group, ipsilateral vs naïve, *P* = 0.0292; contralateral vs naïve, *P* = 0.5075; contralateral vs ipsilateral, *P* = 0.4022; one-way ANOVA-Bonferroni). (d) Left panels show representative blotting band of p-Tau217 and total tau (tau5) in age-matched naïve control group, contralateral hippocampus, and ipsilaterally stimulated hippocampus. Statistical results of p-Tau217 over total tau was quantified and normalized to naïve control (n = 5 per group, ipsilateral vs naïve, *P* = 0.0049; contralateral vs naïve, *P* = 0.0979; contralateral vs ipsilateral, *P* = 0.3905; one-way ANOVA-Bonferroni). Values are mean ± s.e.m.

Consistent with the immunohistochemistry, LTD-inducing LFS significantly enhanced p-Tau181 (**Figure 3c**; **Figure S6**) and p-Tau217 (**Figure 3d**; **Figure S6**) in the ipsilateral hippocampus, while it had no overall significant effect compared with naïve controls on the expression level of p-Tau at the other residues investigated (**Figure S7**). Western blotting indicated that the expression levels of p-Tau181 and p-Tau217, slightly, although not significantly, also increased in the contralateral hippocampus when compared with naïve controls. This indicates that application of LFS activates the commissural pathway sufficiently to enhance the expression levels of both p-Tau181 and p-Tau217 in parts of the contralateral hippocampus.

Enhanced neuronal activity accelerates tauopathy *in vivo* in tau mouse models (Wu *et al*, 2016) and mechanical injury of neurons induces tau mislocalization to dendritic spines (Braun *et al*, 2020). In order to exclude any influence of electrode implantation and baseline recording of evoked field EPSPs on the local expression level of p-Tau, the same experimental processes were performed but no LFS was applied in another cohort of 17-18-month-old rats (**Figure S8a,b**). The immunofluorescent staining results confirmed that no change of p-Tau181 and p-Tau217 levels were present in hippocampus from these rats (**Figure S8c,d**). We thus demonstrate that application of LFS enhances p-Tau181 and p-Tau217 in an age-dependent manner in live rats. The enhancement of tau phosphorylation at both residues by LFS appears more sensitive compared with residues Thr231, Ser202/Thr205, and Ser396.

### Antagonists of NMDA receptors or mGluR5 inhibit the elevation of both p-Tau181 and p-Tau217

Previously, we found that the induction of LTD by LFS, was not blocked by standard systemic doses of either NMDAR or mGluR5 antagonists when applied alone (Hu *et al*., 2014; O’Riordan *et al*., 2018b), while combined systemic injection of both antagonists completely abrogated the induction of synaptic LTD in young adult rats (O’Riordan *et al*., 2018b). Therefore, we wondered if synaptic LTD could be blocked by standard systemic doses of either NMDAR or mGluR5 antagonists when applied alone in relatively aged (17-18-month-old) rats. Similar to our previous finding in young rats (Hu *et al*., 2014; O’Riordan *et al*., 2018b), a standard systemic dose (10 mg/kg, i.p.) of the competitive NMDAR antagonist CPP which completely blocked synaptic long-term potentiation (LTP) in CA3-CA1 synapses (**Figure S9**), did not affect the induction of LTD in aged rats. Thus 1 h post the injection of CPP, the application of LFS suppressed the field EPSPs to 70.2 ± 2.5% in 17-18-month-old rats (**Figure 4a,b**). Next we tested the standard dose of MTEP (3 mg/kg, i.p.) which is known to non-competitively block mGluR5, with a rat hippocampal receptor occupancy of > 90% (Busse *et al*, 2004). The application of LFS induced a depression of field EPSPs to 79.6 ± 3.7% in these MTEP-treated animals (**Figure 4a,b**).

**Figure 4.**
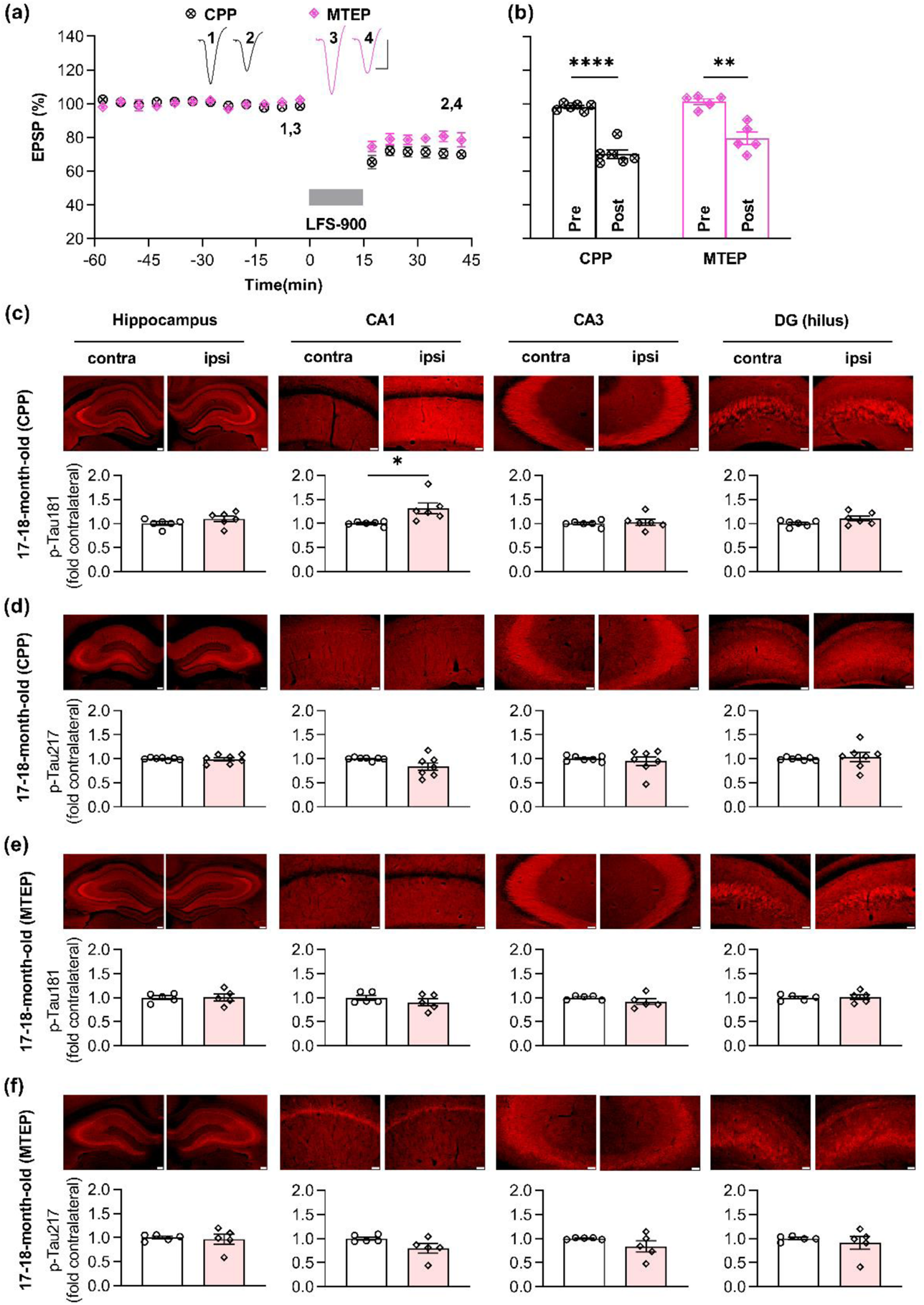
Antagonists of either NMDA receptors or mGluR5 prevent the elevation of both p-Tau181 and p-Tau217 induced by LFS in aged rats. (a) Systemic injection of competitive NMDAR antagonist CPP (10 mg/kg, i.p.) alone 1 h prior to the application of LFS did not affect LTD induction in 17-18-month-old rats. Similarly, intraperitoneal injection of the mGluR5 negative allosteric modulator MTEP (3 mg/kg) alone 1 h did not affect LTD induction by LFS 1 h later in 17-18m old rats. Calibration bars: vertical, 2 mV; horizontal, 10 ms. (b) Summary of the mean EPSP amplitude pre and post-LFS. The EPSP decreased to 70.2 ± 2.5% (n = 6, *P* < 0.0001 compared with pre, paired *t* test) and 79.6 ± 3.7% (n = 5, *P* = 0.0068 compared with pre, paired *t* test) 30 min post-LFS in CPP or MTEP treated rats respectively (two-way RM ANOVA, Treatment *F*_1, 9_ = 6.999, *P* = 0.0267). (c) The upper panel shows the fluorescent images of p-Tau181 labeling (red) and the lower panel displays the corresponding statistical results in CPP-treated 17-18-month-old rats (Scale bar = 200 μm in dorsal hippocampus; scale bars = 50 μm in CA1, CA3 and DG regions). LTD induction by LFS did not affect the level of p-Tau181 in dorsal dorsal hippocampus (*P* = 0.1218), CA3 (*P* = 0.6147), and DG (*P* = 0.0578) except for CA1 region (*P* = 0.0353); paired *t* test. (d) The upper panel shows the fluorescent images of p-Tau217 labeling (red) and the corresponding statistical results are displayed in the lower panel in CPP-treated 17-18-month-old rats. No significant difference was found in p-Tau217 level compared with that in contralateral dorsal hippocampus (*P* = 0.8075), CA1 (*P* = 0.0920), CA3 (*P* = 0.5811), and DG (*P* = 0.7041); paired *t* test. (e) In MTEP-treated aged rats, LFS failed to promote the level of p-Tau181 (red as showed in the upper panel). Summarized mean fluorescence intensities (fold to contralateral) in dorsal hippocampus (*P* = 0.9154), CA1 (*P* = 0.3161), CA3 (*P* = 0.1839), and DG (*P* = 0.8417); paired *t* test. (f) MTEP treatment also prevented the enhancement of p-Tau217 by LFS in aged rats. The upper panel showed the fluorescent images of p-Tau217 (red). The corresponding statistical results were displayed in the lower panel and no significant difference was found in dorsal hippocampus (*P* = 0.7702), CA1 (*P* = 0.1598), CA3 (*P* = 0.2262), and DG (*P* = 0.5311); paired *t* test. Values are mean ± s.e.m.

Given that neither block of NMDAR with CPP nor block of mGluR5 with MTEP prevented the induction of LTD in aged rats, we hypothesized that neither of these treatments on their own would affect the enhancement of p-Tau181 and p-Tau217 by LFS. To test this, we assessed total tau and p-Tau on both residues in the hippocampus of the rats treated with CPP or MTEP. Immunofluorescent staining for antibody Tau46 confirmed that neither agent affected total tau expression level after induction of LTD by LFS in aged rats (**Figure S10**). Surprisingly, even though neither of these treatments on their own affected the induction of LTD, elevation of p-Tau181 and p-Tau217 by LFS were abolished/inhibited in both CPP and MTEP-treated aged rats. After systemic treatment of CPP, the enhancement of p-Tau181 by LFS was strongly inhibited overall although some enhancement was seen in the CA1 area (**Figure 4c**), while the elevation of p-Tau217 by LFS was completely abolished (**Figure 4d**).

Somewhat similarly, the elevation of both p-Tau181 and p-Tau217 triggered by LFS was completely prevented by systemic administration of MTEP (**Figure 4e,f**). CPP is a competitive NMDAR antagonist, so its ability to block NMDARs depends on the magnitude of stimulus-evoked transmitter release during the LFS period. This may help explain why the enhancement of p-Tau181 was resistant to CPP in the CA1 area. Together, these findings indicate that increases of p-Tau181 and p-Tau217 triggered by LFS needs the co-activation of mGluR5 and NMDARs.

### Targeting ageing with a small molecule cognitive enhancer ISRIB blocks the enhancement of p-Tau181 and p-Tau217 by LFS in aged rats

Given that induction of LTD by LFS enhances p-Tau181 and p-Tau217 in an age-dependent manner and ageing is the greatest risk factor for AD (Hou *et al*, 2019), we hypothesized that targeting ageing might prevent the elevation of p-Tau181 and p-Tau217 by LFS. Very recently, Krukowski et al. discovered that the small molecule cognitive enhancer ISRIB reverses age-associated changes in hippocampal neuron function (Krukowski *et al*., 2020). To test this hypothesis, we treated relatively aged (17-18-month-old) rats with the same ISRIB treatment paradigm of daily injections on 3 consecutive days as reported (Krukowski *et al*., 2020), and *in vivo* electrophysiological experiments were carried out on these animals 18 days after the last injection of ISRIB (**Figure 5a**). LFS induced stable and robust LTD in all these rats (**Figure 5b,c**). Immunofluorescent staining for antibody Tau46 confirmed that induction of LTD by LFS did not lead to a change in total tau expression level in ISRIB-treated rats (**Figure S11**). Intriguingly, neither p-Tau181 (**Figure 5d**) nor p-Tau217 (**Figure 5e**) were increased by the same LFS conditioning protocol which triggered enhancement of both p-Tau181 and p-Tau217 in untreated age-matched rats (**Figure 1e** and **Figure 2b**).

**Figure 5.**
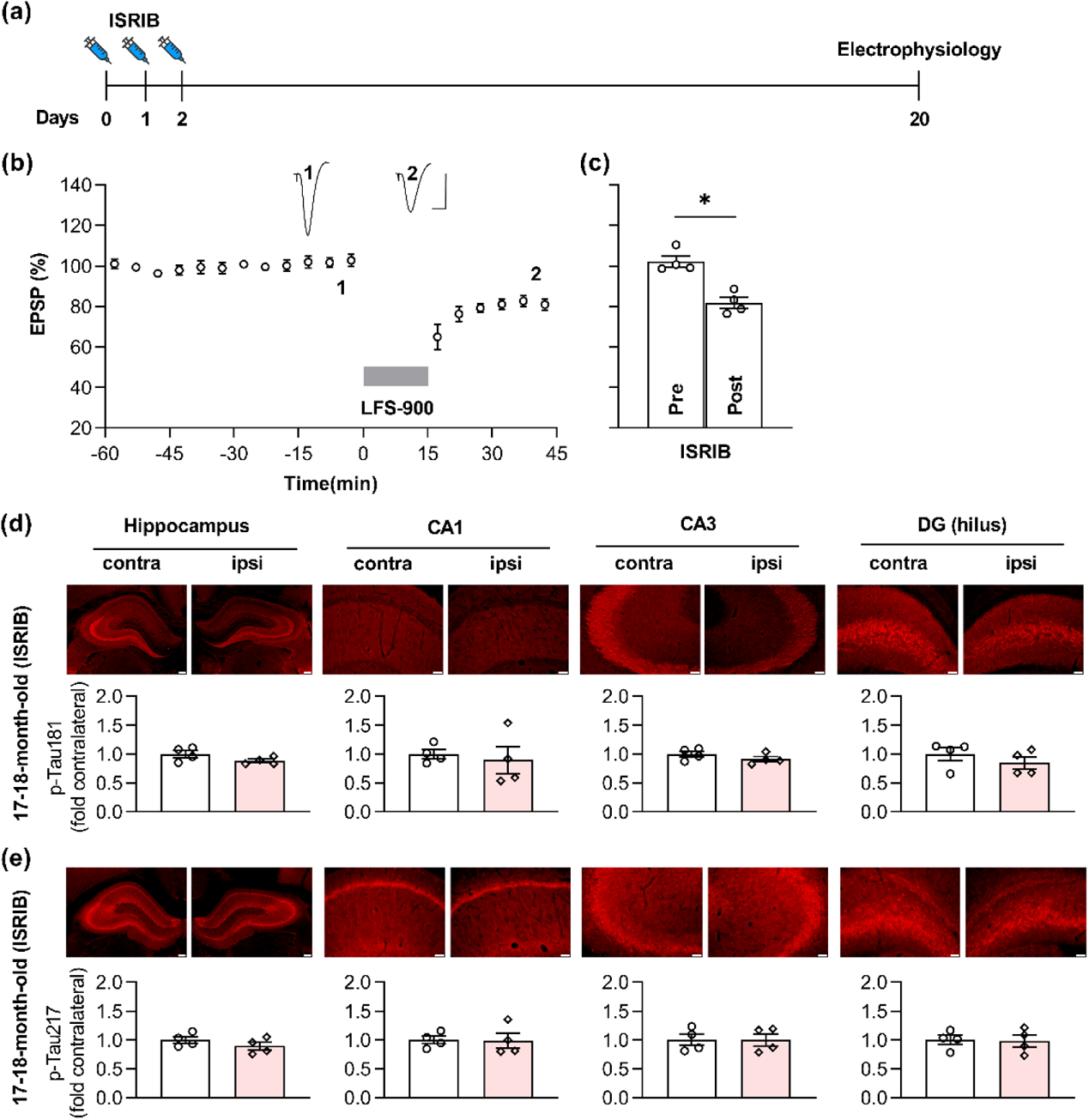
ISRIB blocks LFS-induced enhancement of p-Tau181 and p-Tau217 in aged rats. (a) Experimental paradigm for ISRIB injection and electrophysiology experiments. ISRIB (2.5 mg/kg, i.p.) was systemically injected for 3 days and *in vivo* electrophysiology experiments were performed 18 days after the last injection. (b) LFS induced robust LTD in ISRIB-treated aged rats. Calibration bars: vertical, 2 mV; horizontal, 10 ms. (c) The EPSP decreased to 81.8 ± 2.7% at 30 min post LFS (n = 4, *P* = 0.0211 compared with pre, paired *t* test). (d) Treatment of ISRIB completely prevented the increase of p-Tau181 induced by LFS in aged rats. The upper panel showed the fluorescent images of p-Tau181 labeling (red). As summarized in the lower panel, no significant difference was found in the dorsal hippocampus (*P* = 0.0579), CA1 (*P* = 0.7006), CA3 (*P* = 0.0715), and DG (*P* = 0.2696); paired *t* test. (e) Treatment of ISRIB also successfully blocked the enhancement of p-Tau217 induced by LFS in aged rats. The upper panel showed the fluorescent images of p-Tau217 labeling (red). Summarized statistic results in the lower panel displayed no significant difference in all regions including dorsal hippocampus (*P* = 0.1185), CA1 (*P* = 0.9049), CA3 (*P* = 0.9172), and DG (*P* = 0.6636); paired *t* test. Scale bar = 200 μm in hippocampus, scale bar = 50 μm in CA1, CA3 and DG regions. Values are mean ± s.e.m.

## Discussion

This study reveals that ageing enables the elevation of p-Tau181 and p-Tau217 triggered by LTD-inducing conditioning stimulation in live rats. Growing evidence from many different memory center cohorts indicates that p-Tau181 and p-Tau217 distinguish AD from controls with high accuracy very early in the disease process (Hansson, 2021; Teunissen *et al*, 2022). These findings encourage the prospect that these markers are good enough to become early AD diagnostic tools. Our data provide a novel *in vivo* means to uncover brain plasticity-related cellular and molecular processes of tau phosphorylation at these key sites in health and ageing.

Because LTD induction by LFS at CA3-CA1 synapses in the hippocampus usually necessitates transient elevated synaptic glutamate release and subsequent activation of NMDARs and/or mGluRs (Collingridge *et al*., 2010), we wondered if the enhanced tau phosphorylation triggered by LFS entails these receptors also. Pretreatment with either the NMDAR or mGluR5 antagonist alone completely abrogated the LFS-induced tau phosphorylation, indicating that both receptors need to be activated to trigger enhanced p-Tau181 and p-Tau217. Although we only examined one time point, it is clear that LFS triggers an NMDAR and mGluR5-dependent increase in tau phosphorylation that persists for at least 30 min in all hippocampal subfields examined. Whereas the induction of *in vivo* LTD is only blocked by combined pretreatment with standard doses of CPP and MTEP (O’Riordan *et al*., 2018b), injecting either agent alone (Hu *et al*., 2014; O’Riordan *et al*., 2018b) prevented LFS-triggered tau phosphorylation. In view of these findings, it seems unlikely that enhanced tau phosphorylation at either residues Thr181 or Thr217 is essential for LTD at CA3-CA1 synapses. Also, it remains to be determined if the LFS protocol used in the present study triggers LTD at CA3 recurrent collaterals or back projections as reported previously (Debanne *et al*, 1998). Recent evidence implicates a particular role for extrasynaptic NMDAR in LTD induction (Liu *et al*., 2013; Papouin *et al*., 2012). In the hippocampal CA1 area, mGluR5 is found predominantly located perisynaptically and extrasynaptically on postsynaptic spines of pyramidal cells (Lujan *et al*, 1996; Lujan *et al*, 1997). Co-activation of mGluR5 with NMDARs located in this vicinity is known to strongly enhance the function of the NMDARs, in particular GluN2B-containing NMDARs (Kotecha *et al*, 2003; Sarantis *et al*, 2015). Numerous studies have shown that synaptic NMDARs are mainly involved in normal cognitive function, while extrasynaptic NMDARs are important mediators of neurotoxicity (Vieira *et al*, 2020). Glutamate-induced excitotoxicity increases tau phosphorylation (Sindou *et al*, 1994) and more recent evidence indicates the involvement of extrasynaptic NMDARs activation in tau pathology (Sun *et al*., 2016; Tackenberg *et al*., 2013). Tau is phosphorylated at many different sites by different protein kinases. Both glycogen synthase kinase 3β (GSK3β) and cyclin-dependent kinase-5 (Cdk5) can phosphorylate tau at Thr181 and Thr217 (Liu *et al*, 2002). The activity of GSK3β is significantly enhanced during hippocampal LTD (Peineau *et al*, 2007) but very recent evidence indicates the involvement of GSK3α in both NMDAR-dependent (Draffin *et al*, 2021) and mGluR5-dependent (McCamphill *et al*, 2020) LTD. Cdk5 activation appears to be required for NMDAR-dependent LTD at CA3-CA1 synapses also (Mishiba *et al*, 2014).

Ageing is the primary risk factor for most neurodegenerative diseases including AD (Hou *et al*., 2019). Age-related reduction in glutamate uptake is associated with extrasynaptic NMDAR and mGluR activation at hippocampal CA1 synapses (Potier *et al*, 2010). Very recent evidence indicates that imbalanced synaptic weights undies the aberrant elevated firing characteristics of both CA3 and CA1 pyramidal neurons in aged, learning impaired rats (Buss *et al*., 2021). Intriguingly, Krukowski et al. discovered that a small molecule cognitive enhancer ISRIB reverses the aberrantly elevated integrated stress response (ISR) in aged mice brain and restores age-related changes in hippocampal neuron function (Krukowski *et al*., 2020). In the present study, the same ISRIB treatment paradigm successfully prevented the elevation of both p-Tau181 and p-Tau217 by LFS in aged rats. Although the detailed mechanisms of how ISRIB prevents tau phosphorylation remain to be elucidated, given that ageing is the single strongest risk factor for AD, targeting ageing is likely to provide novel therapeutic avenues for AD (Livingston *et al*, 2020).

Previous reports indicate that p-Tau396 can be enhanced by LFS-900 (1 Hz) in acute hippocampal slices (Kimura *et al*., 2014; Regan *et al*., 2015). In contrast, no change in p-Tau202/205 was triggered by LFS except under the extreme condition of 2 hours LTD induction over a period of 7 days in slice cultures (Taylor *et al*., 2021). The experimental conditions used to detect changes in p-Tau396 in hippocampal slices differ very significantly from our live brain studies, including relatively young age and non-physiological temperature. It is well recognized that p-Tau is developmentally regulated (Bramblett *et al*, 1993; Brion *et al*, 1993; Goedert *et al*, 1993; Hefti *et al*, 2019; Yu *et al*, 2009) and can be hypothermia-induced (Avila & Diaz-Nido, 2004; Bretteville *et al*, 2012; Gratuze *et al*, 2017; Planel *et al*, 2004).

Although not statistically significant, we found slight elevation of p-Tau396 triggered by LFS-900 *in vivo* in this study. Nevertheless, future studies should elucidate if, like the present *in vivo* studies, changes in p-Tau181 and p-Tau217 are more sensitive to LTD-inducing LFS *in vitro*. Phosphorylated tau species have faster tau turnover rates and shorter half-lives in cultured human and rodent neurons compared with that in human and rodent brains (Sato *et al*, 2018). Thus, our findings in live animals provide a valuable experimental model to directly study the generation of p-Tau during learning and memory-related synaptic plasticity and to help develop clinical biomarkers of AD.

Further highlighting that AD is a chronic disease, it is feasible to carry out more detailed time course analysis of phosphorylated tau in our live animal models. To conclude, our data in this study show that, similar to clinical biomarker findings, tau phosphorylation is preferentially triggered at certain sites including Thr181 and Thr217 by LTD-inducing conditioning stimulation in aged brain in live rats. Given current technical and ethical barriers to tau detection in live patient’s brain, our experimental models provide a valuable means to (1) directly study the generation of p-Tau during learning and memory-related synaptic plasticity, (2) help optimize the choice of p-Tau as biomarkers, and (3) aid in the selection of p-Tau directed therapies in future clinical trials of AD.

## Materials and Methods

### Animals

All experiments were performed following the guidelines of the ARRIVE (Animal Research: Reporting of In Vivo Experiments) guidelines 2.0 (Percie du Sert *et al*, 2020) and were approved by the Animal Care and Use Committee of Zhengzhou University, China. All efforts were made to minimize the number of animals used and their suffering.

Young adult (2-3-month-old), and aged (17-18-month-old) male Sprague Dawley rats were provided by the Laboratory Animal Center of Zhengzhou University. The animals were housed under a 12 h light-dark cycle at room temperature (19-22°C) with continuous access to food and water *ad libitum*. Prior to the acute experiments, animals were anaesthetized with urethane (1.5-1.6 g/kg, i.p.). Lignocaine (10 mg, 1% adrenaline, s.c.) was injected over the area of the skull where electrodes and screws were to be implanted. The body temperature of the rats was maintained at 37-38°C with a feedback-controlled heating blanket during the whole period of surgery and recording.

### Electrophysiology

Electrodes were made and implanted as described previously (Hu *et al*., 2014). Briefly, monopolar recording electrodes were constructed from Teflon-coated tungsten wires (75 μm inner core diameter, 112 μm external diameter) and twisted bipolar stimulating electrodes were constructed from Teflon-coated tungsten wires (50 μm inner core diameter, 75 μm external diameter) separately. Field excitatory postsynaptic potentials (EPSPs) were recorded from the stratum radiatum in the CA1 area of left or right hippocampus in response to stimulation of the Schaffer collateral-commissural pathway. Electrode implantation sites were identified using stereotaxic coordinates relative to bregma, with the recording site located 3.4 mm posterior to bregma and 2.5 mm lateral to midline, and stimulating site 4.2 mm posterior to bregma and 3.8 mm lateral to midline. The final placement of electrodes was optimized by using electrophysiological criteria and confirmed via postmortem analysis.

Test EPSPs were evoked by a single square wave pulse (0.2 ms duration) at a frequency of 0.033 Hz and an intensity that triggered a 50% maximum EPSP response. LTD was induced using 1 Hz low frequency stimulation (LFS) consisting of 900 pulses (0.2 ms duration). During the LFS the intensity was raised to trigger EPSPs of 95% maximum amplitude. LTP was induced using 200 Hz high frequency stimulation (HFS) consisting of one set of 10 trains of 20 pulses (inter-train interval of 2 s). The stimulation intensity was raised to trigger EPSPs of 75% maximum during the HFS. None of the conditioning stimulation protocols elicited any detectible abnormal changes in background EEG, which was recorded from the hippocampus throughout the experiments.

### Immunofluorescent staining

After electrophysiological recording under anesthesia of urethane, rats were transcardially perfused with pre-warmed normal saline followed by cold 4% paraformaldehyde in PBS at pH 7.4. Brains were carefully removed and post-fixed in 4% paraformaldehyde for 6 h. Brains were dehydrated in 20% sucrose followed by 30% sucrose. Tissues were rapidly frozen and cut coronally (50 μm). Sections near the stimulating and recording electrodes were reserved and stored in cryoprotective solution (150 mM ethylene, 100 mM glycerol, 250 mM PBS) at −20°C. For total tau and phosphorylated-tau staining, sections were washed three times with PBS for 5 min and then permeabilized with 0.1% Triton X-100 diluted in PBS for 30 min at room temperature. Then sections were blocked in a PBS solution containing 10% normal goat serum, 3% BSA, 0.1% Triton X-100 for 1 h at room temperature. Sections were incubated respectively with the primary antibodies (**Table S1**) (Tau5, Cell Signaling Technology, 46687S, 1:200; Tau46, Cell Signaling Technology, 4019S, 1:200; Phospho-Tau (Thr181), Cell Signaling Technology, 12885S, 1:200; Phospho-Tau (Thr217), ThermoFisher, 44-744, 1:200; Phospho-Tau (Thr231), Abcam, ab151559, 1:200; Phospho-Tau (Ser202, Thr205), ThermoFisher, MN1020, 1:200; Phospho-Tau (Ser396), Affinty, AF3148, 1:200) in a humidified chamber overnight at 4°C. After 24 h, sections were washed three times with PBS for 5 min, followed by incubation with species-specific secondary antibodies conjugated to 488 nm or 568 nm fluorophores (Alexa Fluor® 488, Abcam, 150113, 1:500; Alexa Fluor® 568, Abcam, 175471, 1;500) for 1 h at room temperature. Sections were subsequently washed and stained with DAPI solution (Solarbio, C0065) for 10 min at room temperature. After washing with PBS, sections were mounted on the glass slide with antifade mountant (Southern Biotech, 0100-20) and then processed for imaging using an Olympus fluorescent microscope (BX5WI, Olympus, Japan). ImageJ version 1.52a software (National Institute of Mental Health, Bethesda, Maryland, USA) was used to analyze the intensity of total tau, p-Tau of immunofluorescent staining.

### Western blot

The rats were sacrificed 30 min post-LFS in the LFS-treated group or the same timeline under urethane anaesthesia in the naïve control group. The whole brain was taken out and the hippocampus from both sides was separated. Approximately 2-mm-thick hippocampal tissue surrounding the sites of electrodes was kept and frozen immediately in liquid nitrogen and stored at −80°C. The tissues were homogenized in lysis buffer (10mM Tris-HCl, pH 7.5, 150mM NaCl, and 0.5% Triton X-100, 0.1mM PMSF) containing 1% protease inhibitor Cocktail (Sigma-Aldrich, CW2200S) and 1% phosphatase inhibitor Cocktail (Sigma-Aldrich, CW2383S). The protein concentrations were determined by the BCA Protein Assay Kit (Glpbio, GK10009), and samples were separated by 10% Tris-glycine SDS-PAGE. The proteins were transferred onto polyvinylidene fluoride (PVDF) membranes (Millipore, IPVH00010). Then the membranes were blocked with 5% non-fat milk for 1 h at room temperature. After blocking, the membranes were incubated respectively with the primary antibodies (**Table S1**) overnight at 4°C. After primary antibody incubation, the membranes were washed three times in TBST and then incubated with HRP-conjugated goat anti-rabbit IgG (ZSGB-BIO, ZB-2301, 1:25000) for 2 h at room temperature. Finally, the target protein bands were visualized with chemiluminescence reagents (Shanghai Willget Biotech, F03) and then detected with ProteinSimple System (Hybrid HY8300, FluorChem E system, USA). Quantification of the protein expression was calculated with ImageJ (version 1.52a).

### Pharmacological agents

(R,S)-3-(2-carboxypiperazin-4-yl)propyl-1-phosphonic acid ((±)-CPP, Alomone, C-175) and 3-((2-methyl-1,3-thiazol-4-yl)ethynyl)pyridine hydrochloride (MTEP hydrochloride, Abcam, ab120035) were prepared in distilled water and diluted with saline to the required concentration.

Trans-N,N′-(Cyclohexane-1,4-diyl)bis(2-(4-chlorophenoxy) acetamide (ISRIB, Sigma, SML0843) was dissolved in dimethyl sulfoxide (DMSO) with gentle warming in a 40°C water bath and vortexed until the solution became clear. Then the solution was diluted in polyethylene glycol 400 (PEG400) with gentle warming in a 40°C water bath and vortexed. The solution was prepared freshly and diluted in warm saline (37°C) before injection. 1:1 DMSO and PEG400 in saline was used as vehicle control. The choice of dose and timing of ISRIB administration was based on previous reports (Krukowski *et al*., 2020) and our study of the pharmacokinetics of ISRIB in live rats (Hu *et al*, 2022). Four pair-housed relatively aged (17-18-month-old) rats received a single daily injection of ISRIB (2.5 mg/kg, i.p.) on 3 consecutive days. Then the rats stayed pair-housed for 18 days after the third injection of ISRIB.

### Data analysis

Values are expressed as the mean ± s.e.m. For the electrophysiology experiments, the last 10 min prior to LFS was used to calculate the “Pre”-induction EPSP amplitude. Unless otherwise stated the magnitude of LTD was measured over the last 10 min at the end of recording after (“Post”) LFS. To compare between two group, repeated measures two-way ANOVA with Bonferroni *post hoc* test was used. To compare between groups of three or more, one-way ANOVA with Bonferroni multiple comparisons was used. A two-tailed paired Student’s *t*-test (paired *t*) was used to compare between “Pre” and “Post” within groups. For the immunofluorescence staining, internal comparison was used. Thus, immunofluorescent staining signal was quantified in the entire and selected subregions of the ipsilaterally stimulated hippocampi and the contralateral one from the same animals. The average of the contralateral control was standardized to 1. A two-tailed paired Student’s *t*-test was used to compare between ipsilateral and contralateral. Data of western blotting between conditions were compared using one-way ANOVA with Bonferroni multiple comparisons. A value of *P <* 0.05 was considered statistically significant (\**P* < 0.05, \*\**P* < 0.01, \*\*\**P* < 0.001, \*\*\*\**P* < 0.0001). All data were evaluated and graphed using Prism 9.0 (GraphPad Inc, San Diego, CA, USA).

### Data availability

All raw data supporting the findings of this study are available from the corresponding author upon reasonable request.

## Abbreviations

AD: Alzheimer’s disease
Aβ: amyloid-β protein
Aβ_O_: Aβ oligomers
CSF: cerebrospinal fluid
Cdk5: cyclin-dependent kinase-5
DAPI: 4′,6-diamidino-2-phenylindole
DG: dentate gyrus
DMSO: dimethyl sulfoxide
EPSP: excitatory postsynaptic potential
GSK3α: glycogen synthase kinase 3α
GSK3β: glycogen synthase kinase 3β
HFS: high frequency stimulation
i.c.v.: intracerebroventricular
ISR: integrated stress response
LFS: low frequency stimulation
LTD: long-term depression
LTP: long-term potentiation
mGluR5: metabotropic glutamate receptor subtype 5
NMDA: N-methyl-D-aspartate
NMDAR: N-methyl-D-aspartate receptor
PEG400: polyethylene glycol 400
PMSF: phenylmethylsulfonyl fluoride
PrP^C^: cellular prion protein
PVDF: polyvinylidene fluoride
p-Tau: phosphorylated tau
SDS: sodium dodecyl sulfate
TBS-T: tris-buffered saline containing 0.1% Tween 20

## Acknowledgments

This study has been funded by National Natural Science Foundation of China (U2004134) and Zhengzhou University (140/32310295) to NWH, and by Science Foundation Ireland (19/FFP/6437 and 14/IA/2571) to MJR. The funders had no role in study design, data collection and analysis, decision to publish, or preparation of the manuscript. We thank Professor Seán Kennelly (Tallaght University Hospital, Trinity College Dublin) and Professor Tim Lynch (Mater Misericordiae University Hospital, University College Dublin) for advice.

## Author contributions

NWH and MJR jointly conceptualized the study and NWH directed experiments; YZ, ZH, and PY conducted the electrophysiological experiments; YZ, YY, MZ, and SQ performed immunofluorescent staining; YZ, YY, MZ, and SQ performed Western blot; BL and JX assisted with Western blot and immunofluorescent staining; All authors contributed to preparing figures and data analysis; NWH wrote the first draft of the manuscript; All authors contributed to reviewing and editing the manuscript, and approved its final version.

## Ethics approval

All experiments were performed following the guidelines of the ARRIVE (Animal Research: Reporting of In Vivo Experiments) guidelines 2.0 (Percie du Sert *et al*., 2020) and were approved by the Animal Care and Use Committee of Zhengzhou University, China. All efforts were made to minimize the number of animals used and their suffering.

## Supporting Information Listing

Table S1: Antibodies used in this study

Figure S1-11 and legends

